# Intra-tumor microbiome-based tumor survival indices predict immune interaction and drug sensitivity on pan-cancer scale

**DOI:** 10.1101/2024.12.16.628699

**Authors:** Yan Gao, Haohong Zhang, Dongliang Chu, Kang Ning

**Affiliations:** Key Laboratory of Molecular Biophysics of the Ministry of Education, Hubei Key Laboratory of Bioinformatics and Molecular-imaging, Center of AI Biology, Department of Bioinformatics and Systems Biology, College of Life Science and Technology, Huazhong University of Science and Technology, Wuhan 430074, Hubei, China; Geneis Beijing Co., Ltd., Beijing, 100102, China

## Abstract

Growing research evidence indicates a substantial influence of the intra-tumor microbiome on tumor outcome. However, there is currently no consistent criterion for identifying the association of microbes with tumor progression and response to treatment across various types of cancer. In this study, we concentrate on the intra-tumor microbiome and develop the Tumor Microbiome Survival Index (TMSI), a measure indicative of cancer patient’ survival risk. Our indices revealed notable distinctions between two stratified risk groups for each of the 10 cancer types and could precisely predict patients’ overall survival. For each type of cancer, our findings unveiled two distinct gene expression profiles and shed light on the varying patterns of immune and stromal cell enrichment between the two risk groups. Additionally, we noted that the high-TMSI group exhibited substantially elevated IC50 values for a number of drugs, indicating that individuals in the low-TMSI group might experience superior therapeutic effects from chemotherapy. These findings illuminate the complex dynamics between the tumor microbiome, the patient’s immune reaction, and medical outcomes, thus shedding light on microbiome-based personalized therapeutic interventions.

**Highlights:**

1. The TMSI model generates a survival risk score based on the intra-tumor microbiome.
2. The TMSI model predicts survival rates accurately across multiple cancer types.
3. Molecular-level analysis highlights distinct immune interactions with tumor microbes.
4. High-TMSI groups showed altered drug sensitivity and implying potential differences in treatment response.

## Introduction

Cancer is the second-leading cause of death worldwide, accounting for approximately 2.6 million deaths in China, 2022 [1, 2]. Traditionally, cancer has been regarded as a disease originating from alterations in the genetic makeup of human beings [3, 4]. Consequently, there has long been a recognized connection between tumor gene expression and cancer outcomes [5–8]. However, while genetic markers have been instrumental in advancing our understanding of cancer progression, their clinical applicability is limited. Relying solely on genetic markers provides an incomplete comprehension of cancer biology [9].

A tumor is not merely a cluster of cancer cells; it is a heterogeneous assembly of infiltrating and resident host cells, secreted factors, extracellular matrix, and possibly microbes, all of which influence prognosis [10]. This complexity is further compounded by the genetic heterogeneity that tumors often exhibit, even within the same cancer type. Such heterogeneity arises from factors including mutations, chromosomal rearrangements, and epigenetic modifications [9]. This heterogeneity can lead to variations in gene expression patterns among different tumors and even within different regions of the same tumor. Therefore, integrating multi-omics data is essential for prognostic analysis [11, 12].

The tumor microbiome, a complex and diverse community of microbes that inhabit human tumors and adjacent tissue, has gained increasing attention for its potential role in cancer biology. [13, 14]. Although there is still debate regarding the extent of the relationship between microbes and cancer, a growing body of evidence supports the involvement of the tumor microbiome in various types of cancer. Poore et al. [15] recently developed a computational workflow that utilizes two orthogonal microbial detection pipelines to obtain high quality microbial abundances from high-throughput sequencing data of human tumors. The tumor microbiome has been shown to play certain functions in the development [16], progression [17], and response to treatment [18] for various types of cancer. The relationship between the tumor microbiome and patient survival is the subject of ongoing research, as some studies [19, 20] have suggested that the microbiome of a tumor may impact patient survival. For example, Malassezia globose has been linked to a higher risk of mortality in breast cancer patients, underscoring the potential impact of specific microbes on patient outcomes [20]. Therefore, the intra-tumor microbial signal could serve as a potential candidate for accurately prognostic prediction. Nevertheless, a consistent criterion for capturing these survival-related microbes in different tumors is lacking, and the mechanisms through which microbiota contribute to shaping the molecular properties of tumors and influencing clinical outcomes remain poorly understood [21–23].

Given the growing recognition of the tumor microbiome’s role in cancer prognosis, we introduce the Tumor Microbiome Survival Index (TMSI), a survival risk score based solely on the tumor microbiome. Applied to a cohort of cancer microbiome dataset from Emile Voest et al. [24], our index demonstrated significant differentiation of two risk groups in 10 cancer types and accurately predicted the 1-year, 2-year, and 4-year overall survival (OS) rates. The TMSI has also been validated on an external pancreatic cancer dataset [25]. We then analyzed the molecular-level differences between the two risk groups using paired gene expression and DNA methylation data, focusing on potential crosstalk between the host immune system and the tumor microbiota. Our findings revealed distinct patterns of immune cell and stromal cell enrichment between the two risk groups, with certain microbiota potentially regulating immune gene expression by influencing DNA methylation levels. Furthermore, we observed significantly higher IC50 values for several drugs in the high-risk group, suggesting that patients in the low-risk group may derive greater benefit from these medications. These results shed light on the intricate interplay between the tumor microbiome, host immune response, and clinical outcomes, thereby offering new avenues for personalized therapeutic interventions.

## Methods

### Construction of the TMSI Model

Utilizing microbiome data from tumor tissues of 3,052 samples across 10 types of cancer, TMSI model was constructed. Initially, the "survival" R package (version 3.6-4) was employed to conduct univariate analysis through the proportional hazards model, and bacteria with P-values less than 0.05 were selected as prognostic-related bacteria. Subsequently, the random survival forest (RSF) model from the "randomForestSRC" R package (version 3.3.0) was used to further filter candidate bacteria closely related to survival.

The parameters for the random survival forest analysis are specified as follows: the number of trees, ntree, is set to 2000; the minimum node size, nodesize, is 20; the dataset used is rf_data; the number of splits, nsplit, is 20. The method for handling missing data is set to na.impute, and tree-level error estimation is enabled by tree.err. The importance of variables will be assessed as importance is set to TRUE. The random seed for reproducibility is set to 999.

The algorithm ranked each bacterium according to importance, and we selected the 100 most important bacteria for subsequent analysis. Through the "glmnet" R package (version 4.1-8), these 100 bacteria were used as inputs for the least absolute shrinkage and selection operator (LASSO) Cox regression model, ultimately screening out the significant bacteria. Based on the abundance of the corresponding bacteria and the Cox coefficients of the patients, the Tumor Microenvironment-based Immune Score Indices (TMSIs) for each patient can be calculated using the algorithm of the inner product of matrices. The calculation method has been publicly announced as follows:

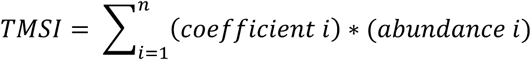

In this formula, *n* represents the total number of independent prognostic tumor microbiome biomarkers selected by univariate and multivariate Cox regression for predicting patient survival. The "*coefficient i*" refers to the regression coefficient of the i-th tumor microbiome biomarker in the multivariate Cox regression, and "*abundance i*" refers to the microbial abundance of the i-th tumor microbiome biomarker.

We found that tumor microbiome prognostic markers independently affect patients’ OS in all cancers. Utilizing the optimal cutoff value determined by the "surv_cutpoint" function, patients within these 10 cancer types were stratified into TMSI-high and TMSI-low groups. To evaluate the predictive performance of TMSI in estimating patients’ OS, Kaplan-Meier curves and ROC curves were employed.

### Evaluation and Validation of the TMSI Model

In order to improve the forecasting accuracy of 1-year, 2-year, and 4-year survival rates across various cancers, we amalgamated clinical characteristics (including age, subtype, gender, and stage) with TMSI to develop a prognostic nomogram model. This was accomplished utilizing the "rms" (v6.8.0) package. The performance of the model was assessed using time-dependent Receiver Operating Characteristic (ROC) curves and the concordance index (C-index).

In our study of the cohort of 10 types of cancer, we specifically validated pancreatic adenocarcinoma (PAAD) using the PRJNA542615 dataset. This dataset includes 16S rRNA sequencing data from 43 tumor tissue samples with known survival status. In the discovery cohort, we identified 10 genera of bacteria to construct the TMSI model, but four of them were not detected in the validation cohort. To address this issue, we employed an interpolation method, using the average abundance of these bacteria from the discovery cohort as the estimated abundance for the corresponding bacteria in each sample of the validation cohort. Subsequently, we used the bacterial coefficients determined from the discovery cohort to calculate the TMSI for each sample in the validation cohort.

### Integrated analysis of the tumor microenvironment and the tumor microbiome

To describe the tumor microenvironment landscape of patients in TMSI-high and TMSI-low groups, matching tumor RNA-seq data of these two subtypes across cancers was introduced. The differentially expressed genes (DEGs) between these two groups were calculated by DESeq2 (v 1.44.0) package using the raw count of RNA-seq with the thresholds (|log2FoldChange| < 1 and adjust P-value < 0.05). Gene lists for Kyoto Encyclopedia of Genes and Genomes (KEGG) and Gene Ontology (GO) were sourced from the MSigDB database (https://www.gsea-msigdb.org/gsea/msigdb/). Subsequently, Gene Set Enrichment Analysis (GSEA) was conducted utilizing the "clusterProfiler" (v4.10.1) package. Immune-related DEGs were annotated and mapped to GO terms and KEGG pathways using the ImmPort database (https://www.immport.org/home). The prognostic significance of these immune-related DEGs was evaluated through univariate Cox regression analysis and Kaplan-Meier survival curves. Furthermore, we explored the underlying correlation between immune-related DEGs and circulating microbial prognostic markers. Additionally, immune cell infiltration in tumors was assessed using "xCell" (v1.1.0) package. Finally, expression levels of immune checkpoint molecules such as PD-1/PD-L1 and CTLA4 were compared between the two groups.

### Mediation Analysis

We employed the mediation (v4.5.0) package to perform mediation analysis, with the alteration in microbial abundance as the independent variable (X), the change in gene expression as the dependent variable (Y), and the change in gene methylation as the potential mediator variable (M). The beta value indicates the strength of the influence of the independent variable (X) on the mediator variable (M) and the mediator variable (M) on the dependent variable (Y). Specifically, the beta value is a regression coefficient that reflects the linear relationship and its direction between the independent and mediator variables, as well as between the mediator and dependent variables. In standardized regression, the beta value represents the correlation between the independent and dependent variables. Standardized regression coefficients facilitate comparisons across different units or variables by eliminating the impact of different variable scales. Moreover, when the beta value is statistically significant (indicated by a p-value less than 0.05), it signifies that the independent variable can effectively predict variations in the dependent variable, implying a significant impact of the independent variable on the dependent variable.

### Drug Sensitivity

We set out to evaluate the predictive significance of interactions between the tumor microenvironment and the immune system in forecasting the success of immunotherapy and chemotherapy treatments. The Tumor Immune Dysfunction and Exclusion (TIDE) score, accessible at http://tide.dfci.harvard.edu/, served as the tool for predicting immunotherapy response across various cancer types. Furthermore, to ascertain the sensitivity of patients to chemotherapy drugs, we leveraged the "oncoPredict" package (v1.2) in R. This tool facilitated the computation of the half-maximal inhibitory concentration (IC50) for each patient, thereby enabling us to predict their likely response to chemotherapy agents.

### Statistical Analysis

All statistical analyses and graphical representations were conducted using R software (R version 4.3.1). The Wilcoxon rank-sum test served to compare continuous variables between the TMSI-high and TMSI-low groups. A significance threshold of p < 0.05 was adopted to ascertain statistical significance.

## Results

### Cohorts and Workflow

Our discovery cohort encompasses 3,052 tumor tissue samples across 10 cancer types, with paired omics data comprising microbiome profile, RNA-seq, and methylation array data. Microbiome profile data were obtained from Emile Voest et al. [24], subjected to Kraken2 and PathSeq microbial profiling pipeline. After rigorous decontamination to eliminate potential contaminants, the dataset was refined to 1041 microbial genera. RNA-seq and methylation array data were sourced from The Cancer Genome Atlas (TCGA) via the Genomic Data Commons (https://portal.gdc.cancer.gov/). The meta-information accompanying these datasets includes clinical parameters such as age, gender, and cancer stage, as well as survival data encompassing survival status and time. For validation, we used the PRJNA542615 PAAD dataset as validation cohort, which includes 16S rRNA sequencing data from 43 tumor tissue samples with known survival status [25].

For the association analysis and comparison, we initially identified tumor microbes with independent prognostic significance using univariate cox regression, RSF model and LASSO-Cox regression. The TMSI score for each patient was then calculated by combining the abundance of OS-related microbes and the coefficients from the LASSO-Cox regression. Based on the their TMSI scores, patients were divided into TMSI-high and TMSI-low groups across each type of cancer. To explore the molecular underpinnings of TMSI score variation, we incorporated the gene expression data and methylation data from the host’s tumor tissue and examined the tumor gene expression profiles to investigate their relationships with TMSI scores. Ultimately, we evaluated the TMSI score’s capability to predict responses to cancer therapies, emphasizing its potential to forecast the success of chemotherapy.

### Construction of the TMSI Model

Firstly, we employed univariate Cox regression analysis to identify genera significantly associated with the OS of cancer patients across 10 cancer types (**Figure 2A**). This analysis revealed both risk factors, with a hazard ratio (HR) > 1(P < 0.05), and protective factors, with HR < 1 (P < 0.05). Subsequently, we utilized the random survival forest model to select the 100 most important bacteria and further performed LASSO-Cox regression analysis to delineate genera that independently impact OS (**Figure 2B**). In all 10 types of cancer, we successfully identified genera that independently affect OS. These results formed the basis of the TMSI model, which calculates a patient’s mortality risk by multiplying the abundance of OS-related genera with the coefficients derived from the LASSO-Cox regression. The optimal cutoff value for the TMSI was determined using the "surv_cutpoint" function, thereby stratifying patients into high-TMSI and low-TMSI groups. The Kaplan-Meier curve and the Log-rank test revealed that patients in the high-TMSI group had shorter OS compared to those in the low-TMSI group (**Figure 2G, 2K** and **Supplementary Figure 1 A-I**), with the number of samples in each group and the p-value of the Log-rank test displayed in **Figure 2C**.

**Fig 1.**
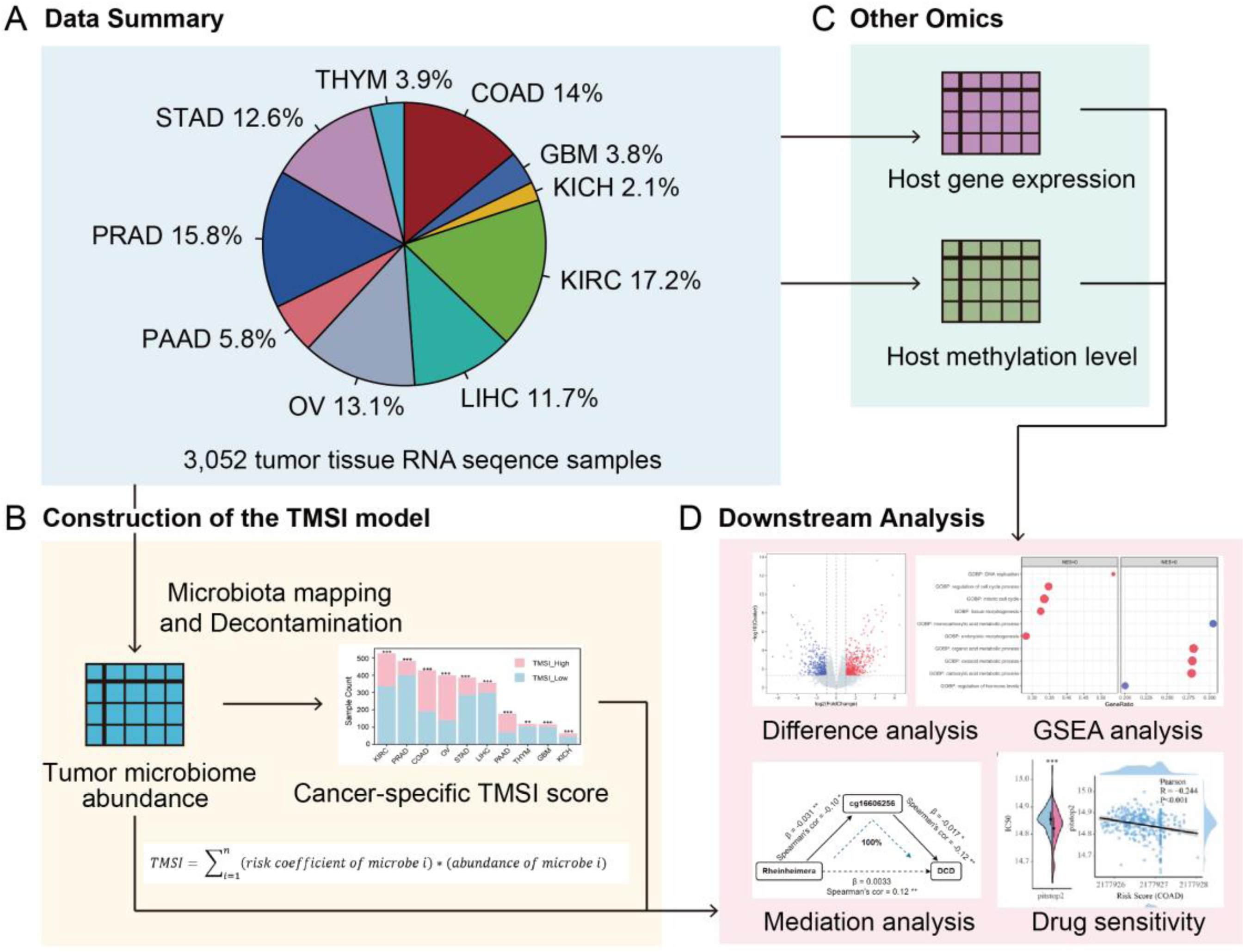
The workflow for a comprehensive analysis of the tumor microbiome across various cancer types. **A**. Data summary. Our discovery cohort includes 3,052 tumor tissue samples across 10 cancer types. **B**. Development of the TMSI model based on tumor microbiome profile data. **C**. Integration of hosts’ RNA-seq data and methylation array data. **D**. Subsequent downstream analysis.

**Fig 2.**
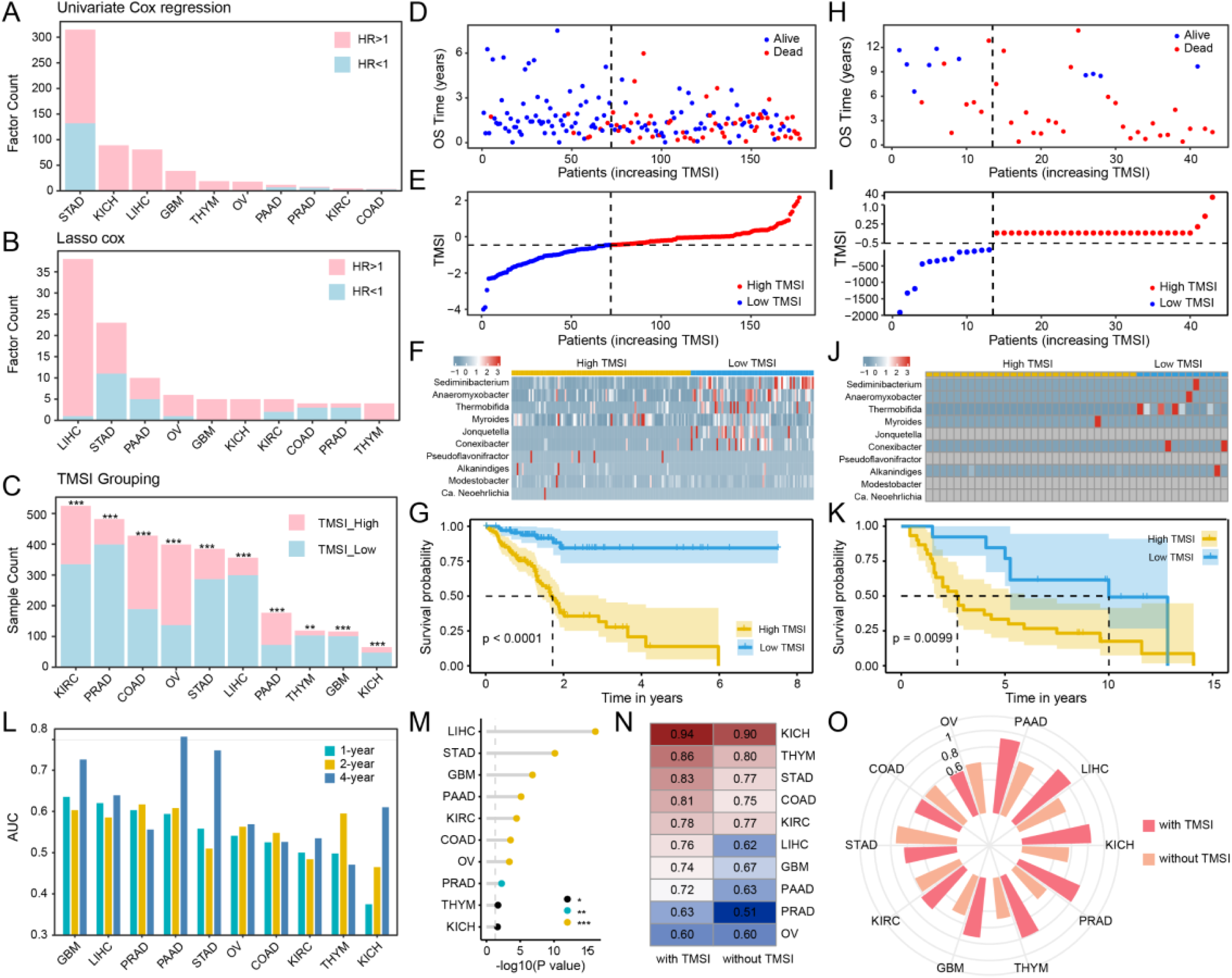
Construction, evaluation and validation of TMSI model to predict the prognosis of patients with ten types of cancer. **A**. Bar graph showing candidate microbial factors significantly associated with OS by univariate Cox regression analysis. **B**. Bar graph of microbial factors independently impacting OS by Lasso-Cox regression analysis. **C**. Significance markers above the bars display the p-values calculated by the Log-rank test between the TMSI-high and TMSI-low groups. **D-G:** Performance of the TMSI model for PAAD in the discovery cohort, **H-K:** Performance in the PAAD validation set. **D, H:** Each point in the scatter plot represents the survival status and survival time of a patient. The horizontal coordinates are the patients ranked from lowest to highest according to their TMSIs. **E, I:** Based on the risk score of each point in the scatter plot representing one patient, they are divided into TMSI-high and TMSI-low groups. **F, J:** The heatmap demonstrates the expression levels of the ten TMSI model bacteria in patients. **G, K:** The Kaplan-Meier curves show the OS time of the two risk groups of patients. **L**. The Area Under the Curve (AUC) of the Receiver Operating Characteristic (ROC) curve shows the performance of TMSI in predicting the 1-year, 2-year, and 4-year OS rates. **M**. The lollipop chart illustrates whether TMSI has an independent impact on prognosis after adjusting for clinical characteristics (including age, subtype, gender, and stage). **N**. The C-index value of the nomogram model based on clinical factors with or without the inclusion of TMSI. **O**. The AUC values of the time-dependent ROC curves show the performance of the nomogram models with or without the inclusion of TMSI in predicting the 4-year OS probabilities of the patients.

### Validation and Evaluation of the TMSI Model

To validate the TMSI model on an external cohort, we used the PRJNA542615 dataset, which includes 16S rRNA sequencing data from 43 tumor tissue samples. In the discovery cohort, we identified 10 genera of bacteria to construct the TMSI model, but four of them were not detected in the validation cohort. To address this issue, we employed an interpolation method, using the average abundance of these bacteria from the discovery cohort as the estimated abundance for the corresponding bacteria in each sample of the validation cohort. Subsequently, we used the PAAD TMSI formula from the discovery cohort to calculate the TMSI for each sample in the PAAD validation set. Using the TMSI formula established from the PAAD samples in the discovery cohort, the TMSI was calculated for each sample in the PAAD discovery cohort (**Figure 2D**) and for each sample in the PAAD validation set (**Figure 2H**). Based on the optimal cutoff value determined from the PAAD dataset in the discovery cohort, both the PAAD dataset in the discovery cohort (**Figure 2E**) and the PAAD validation set (**Figure 2I**) were divided into low TMSI and high TMSI groups. Excluding the four bacteria not detected in the validation set, the other bacteria used to construct the PAAD TMSI had a similar abundance distribution in the discovery cohort (**Figure 2F**) and the validation set (**Figure 2J**). Additionally, Kaplan-Meier analysis showed that patients with a higher TMSI had worse OS in both the discovery cohort (**Figure 2F**) and the validation set (**Figure 2J**).

The TMSI exhibited robust performance in predicting 1-year, 2-year, and 4-year OS rates. Specifically, the AUC values for predicting the 1-year OS rates for GBM and LIHC were 0.635 and 0.620, respectively. Moreover, the AUC values for predicting the 4-year OS rates for PAAD, STAD, and GBM were 0.781, 0.748, and 0.726, respectively **(Figure 2L)**. To ascertain whether the TMSI could function as an independent prognostic indicator, we conducted a multivariate Cox regression analysis based on the TMSI and clinical characteristics, including age, subtype, gender, and stage. The analysis results demonstrated that the TMSI independently affected OS in ten cancers **(Figure 2M)**.

Subsequently, we formulated nomogram survival models for these cancers integrating clinical factors, both with and without the TMSI, to predict the 1-year, 2-year, and 4-year OS probabilities for patients. The nomogram models incorporating the TMSI exhibited higher C-index values compared to those without the TMSI **(Figure 2N)**. The time-dependent ROC curves in predicting the 1-year (**Supplementary Figure 1 J**), 2-year (**Supplementary Figure 1 K**) and 4-year **(Figure 2O)** OS probabilities of the patients suggest that the TMSI enhanced the accuracy of prognostic prediction.

In summary, we have identified microbial genera significantly associated with the prognosis of cancer patients and validated the effectiveness of the TMSI model constructed from these genera in predicting prognosis. The TMSI shows promise as a valuable tool for enhancing the accuracy of prognostic models.

### The stratification of TMSI score exhibits distinct patterns of gene expression

Then we examined the tumor gene expression profiles and investigated their relationships with TMSI scores. We discerned differentially expressed genes (DEGs) (adjusted P-value < 0.05 and |log2 fold change| > 1) between TMSI-high and TMSI-low groups. Across 10 cancers, an asymmetrical distribution in the count of upregulated and downregulated DEGs was evident **(Figure 3A)**. Gene Set Enrichment Analysis (GSEA) outcomes consistently revealed the enrichment of canonical proliferative gene sets, such as "Cell Cycle", "DNA Replication" and "Mitotic Cell Cycle Process" in the TMSI-high group. However, we only observed immune-related gene sets such as "Defense Response to Bacterium" and "Antibacterial Humoral Response" exhibited predominant enrichment in the low-TSMI group in STAD, THYM and COAD **(Figure 3B)**.

**Fig 3.**
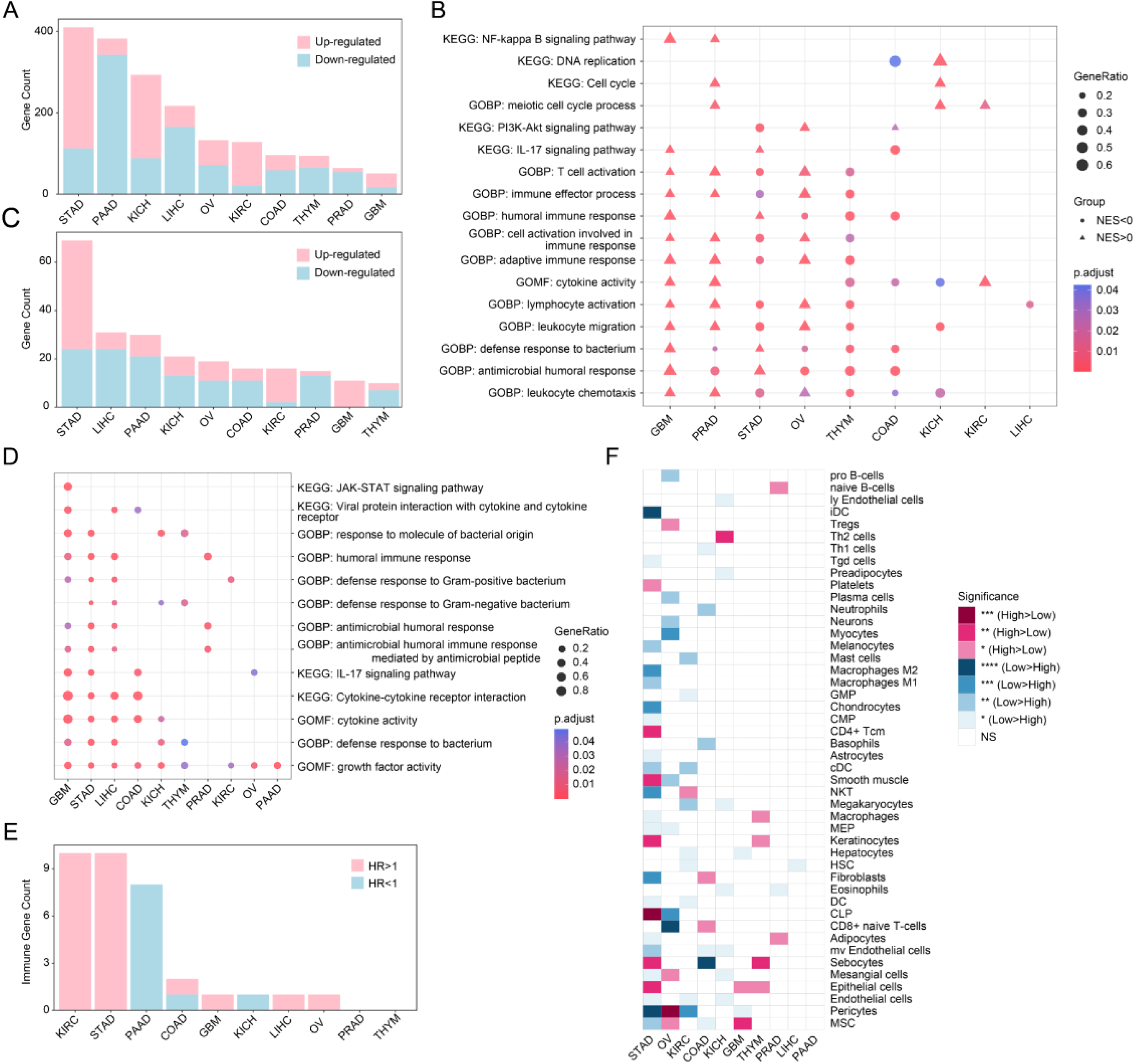
Elucidation of the tumor immune microenvironment through TMSI model. **A**. Bar chart summarizes the differentially expressed genes (DEGs) (P.adj < 0.05 and |log2 FC| > 1) in tumor tissues between the TMSI-high group and the low-TMSI group as determined by DESeq2 analysis. **B**. Bubble chart displays the frequently occurring KEGG pathways and GO terms identified by GSEA enrichment analysis. **C**. Bar chart tallies the immune-related DEGs in tumors within the TMSI-high group compared to the TMSI-low group. **D**. GO and KEGG enrichment analysis based on the immune-related DEGs within each cancer type. **E**. The number of survival-related and immune-related DEGs identified through univariate cox regression and log-rank test across each type of cancer. **F**. Differences in xCell scores of immune cells and stromal cells between TMSI-high group and TMSI-low group. (**F**, ****, p<0.0001; ***, p<0.001; **, p<0.01; *, p<0.05)

Interestingly, certain immune gene sets such as "Lymphocyte Activation" and "Cell Activation Involved in Immune Response," alongside immune pathways like the "IL-17 Signaling Pathway," demonstrated enrichment in the TMSI-high group in GBM, PRAD and OV. This observation may stem from an immune-suppressive state within the corresponding tumor microenvironment, wherein tumor cells exert control over immune cells. Despite apparent gene expression activity, the resultant effect tends toward immune suppression rather than activation. These findings underscore the intricate interplay between tumor TMSIs and the tumor immune microenvironment. High TMSI scores may correlate with increased immune suppression and cell proliferation, while low TMSI scores may be associated with enhanced immune activation and engagement of metabolic pathways. Understanding these mechanisms may provide insights into immune regulation within tumors and support the development of novel therapeutic strategies.

To further explore immune-related DEGs, we used data from the ImmPort database to identify immune-associated Differentially Expressed Genes (DEGs) (P.adj < 0.05 and |log2 FC| > 1) within tumor tissues, distinguishing between the TMSI-high and TMSI-low cohorts. The distribution of upregulated and downregulated immune-related DEGs across various cancers paralleled that of the overall DEGs (depicted in **Figure 3A** and **3C**). Additionally, Gene Ontology (GO) and Kyoto Encyclopedia of Genes and Genomes (KEGG) enrichment analyses revealed the involvement of these immune-related DEGs in “Humoral Immune Response”, “Defense Response to Bacterium”, “Cytokine-Cytokine Receptor Interaction”, and signaling pathways intimately linked to immune system function **(Figure 3D)**. Further analysis involved identifying immune-related DEGs significantly correlated with overall cancer patient survival via univariate Cox regression and Kaplan-Meier survival analysis **(Figure 3E)**. In the histogram, pink bars represent immune-related DEGs deemed risk factors for survival (hazard ratio (HR) > 1; P < 0.05), while blue bars signify those considered favorable factors (HR < 1; P < 0.05). Moreover, tumor immune infiltration analysis unveiled varying degrees of immune cell infiltration between tumors with high and low TMSIs **(Figure 3F)**. Tumors with high TMSI scores exhibited increased immune surveillance, suggesting stronger immune responses. However, this may also implies potential resistance to immune attacks through mechanisms such as upregulation of immune checkpoint molecules, contributing to a higher risk of poor survival outcomes.

### The regulation of host gene expression by tumor microbiome is mediated by host gene methylation

We conducted an in-depth exploration of the dynamic interplay between the host’s immune system and microbial inhabitants, scrutinizing the mechanisms by which the tumor microbiota might modulate gene expression through the alteration of gene methylation patterns. We observed a positive correlation between prognosis risk microbes with prognosis risk genes, while prognosis favorable microbes exhibited a negative correlation with prognosis risk genes in PAAD, COAD and KIRC, and the converse relationship holds true as well **(Figure 4A-C)**. Following the correlation analysis between genes and microbes, we conducted further investigations into the role of methylation in the regulation of host gene expression by microbes. Employing a mediation analysis approach, we aimed to evaluate whether tumor microbes could indirectly regulate the expression of specific genes by influencing their methylation status. In PAAD, *Sediminibacterium* promoted the expression of FGF17 by enhancing methylation at the cg02302670 site, where methylation typically promotes the expression of FGF17 **(Figure 4D)**. These findings imply that microbes may exert modulation over gene expression by altering methylation patterns, thus potentially influencing tumor progression. This discovery sheds light on the intricate mechanisms by which the tumor microbiome interacts with the host’s genetic and epigenetic landscape, offering new insights into the role of microbial communities in cancer development and progression.

**Fig 4.**
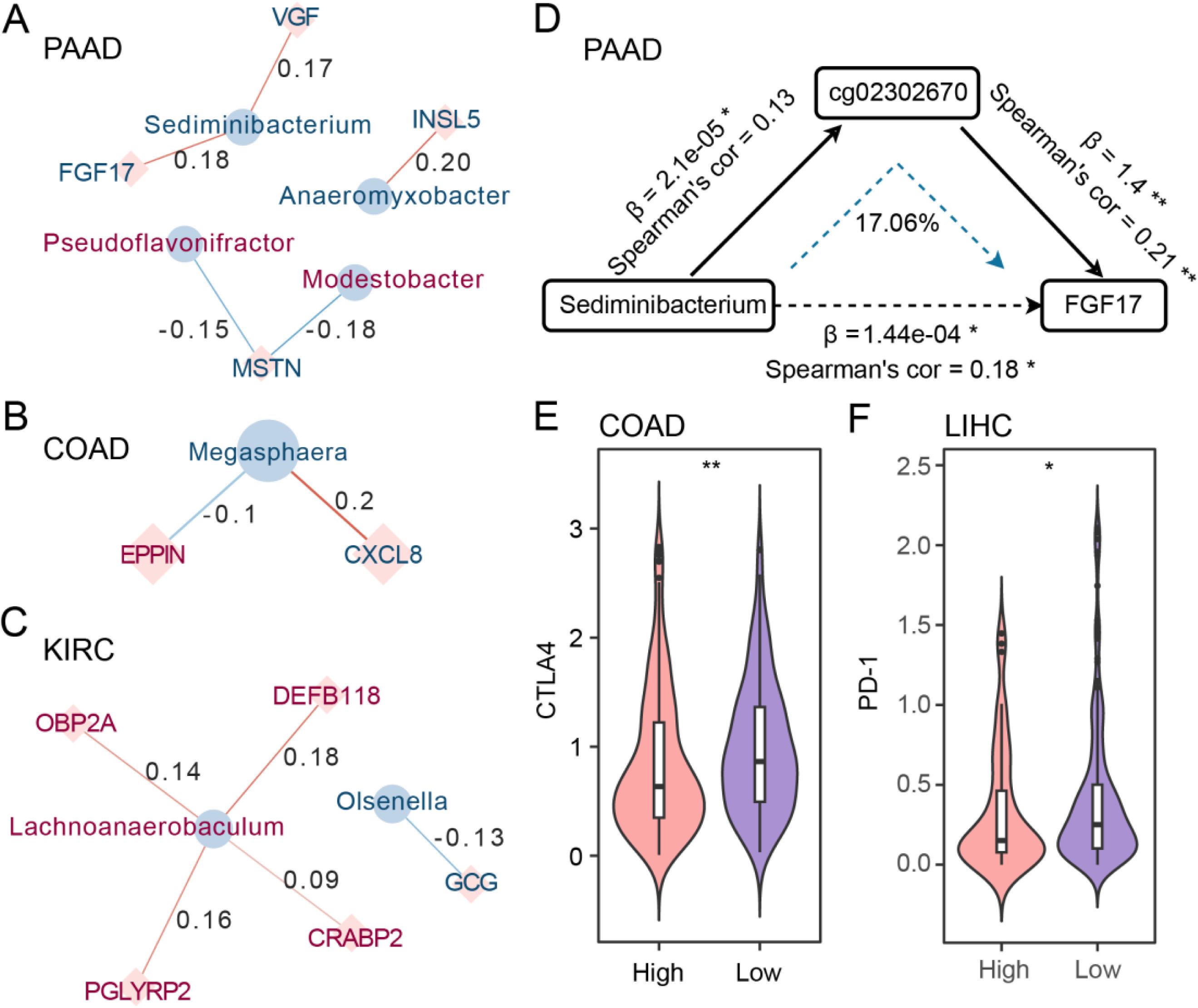
Multi-omics Mediation Effect Analysis. **A-C**. Correlation network analysis of the relationship between model microbes and OS-related immune DEGs of **A** PAAD, **B** COAD, **C** KIRC. Blue circles represent TMSI microbes, and red squares represent immune-related genes associated with the survival of cancer patients. The red text indicates microbes/genes that are beneficial to patient survival, while the blue text indicates microbes/genes that are detrimental to patient survival. **D**. Triangular diagram of mediator effects among model microbe, gene methylation, and immune-related gene in PAAD. **E-F**. CTLA4 expression levels in COAD **E** and PD-1 expression levels in LIHC **F** between TMSI-high and TMSI-low groups (**, p<0.01; *, p<0.05).

### TMSI Predicts the Chemotherapy Efficacy

Microbes play a crucial role in modulating the responses of patients to cancer therapeutics, underscoring their significant influence on treatment efficacy. Numerous studies have affirmed that patients with elevated expression levels of PD-1 or CTLA4 tend to derive greater benefits from immunotherapy. In our study, we observed significant differences in the expression of the immune checkpoint molecule CTLA4 between TMSI-high and TMSI-low groups in COAD **(Figure 4E)**, and similarly, significant differences in the expression of the immune checkpoint molecule PD-1 between TMSI-high and TMSI-low groups in LIHC **(Figure 4F)**. In both cases, the expression was higher in the TMSI-low group compared to the TMSI-high group, suggesting that patients in the TMSI-high group may have a higher risk of survival because they are less likely to benefit from immunotherapy. Subsequently, we conducted TIDE (Tumor Immune Dysfunction and Exclusion) immunotherapy response assessments to explore the predictive value of TMSI for the outcomes of immunotherapy. In 10 types of cancer, no significant differences were found between the TMSI-high and TMSI-low groups, indicating that TMSI has limited capability in predicting the efficacy of immunotherapy **(Supplementary Figure 2)**.

To further elucidate the relationship between TMSI and sensitivity to chemotherapy drugs, we employed the "oncoPredict" software package to forecast the IC50 values for each drug across different cancers. Substantial disparities in chemotherapy responses were evident between high and low TMSI groups across 10 cancers, with STAD displaying the most drugs with significant differences **(Figure 5A)**. Among the top 50 high-frequency drugs significantly correlated with TMSI in each cancer type, a greater number of drugs exhibited IC50 values significantly higher in the high TMSI group compared to the low TMSI group **(Figure 5B)**. Specifically, IC50 values of SGX−523 in KICH **(Figure 5C)**, Procarbazine in KICH **(Figure 5D)**, MI−2 in STAD **(Figure 5E)**, Daporinad in THYM **(Figure 5F)**, BRD−K34222889 in PAAD **(Figure 5G)** and LY−2157299 in PAAD **(Figure 5H)**, were significantly higher in the high TMSI group compared to the low TMSI group, indicating higher resistance to these chemotherapy drugs among patients in the high TMSI group. Conversely, IC50 values of AM−580 in KICH **(Figure 5I)** and NSC632839 in KICH **(Figure 5J)** were significantly higher in the low TMSI group compared to the high TMSI group, suggesting potential benefits for patients in the high TMSI group from these drugs **(Supplementary Table 1)**.

**Fig 5.**
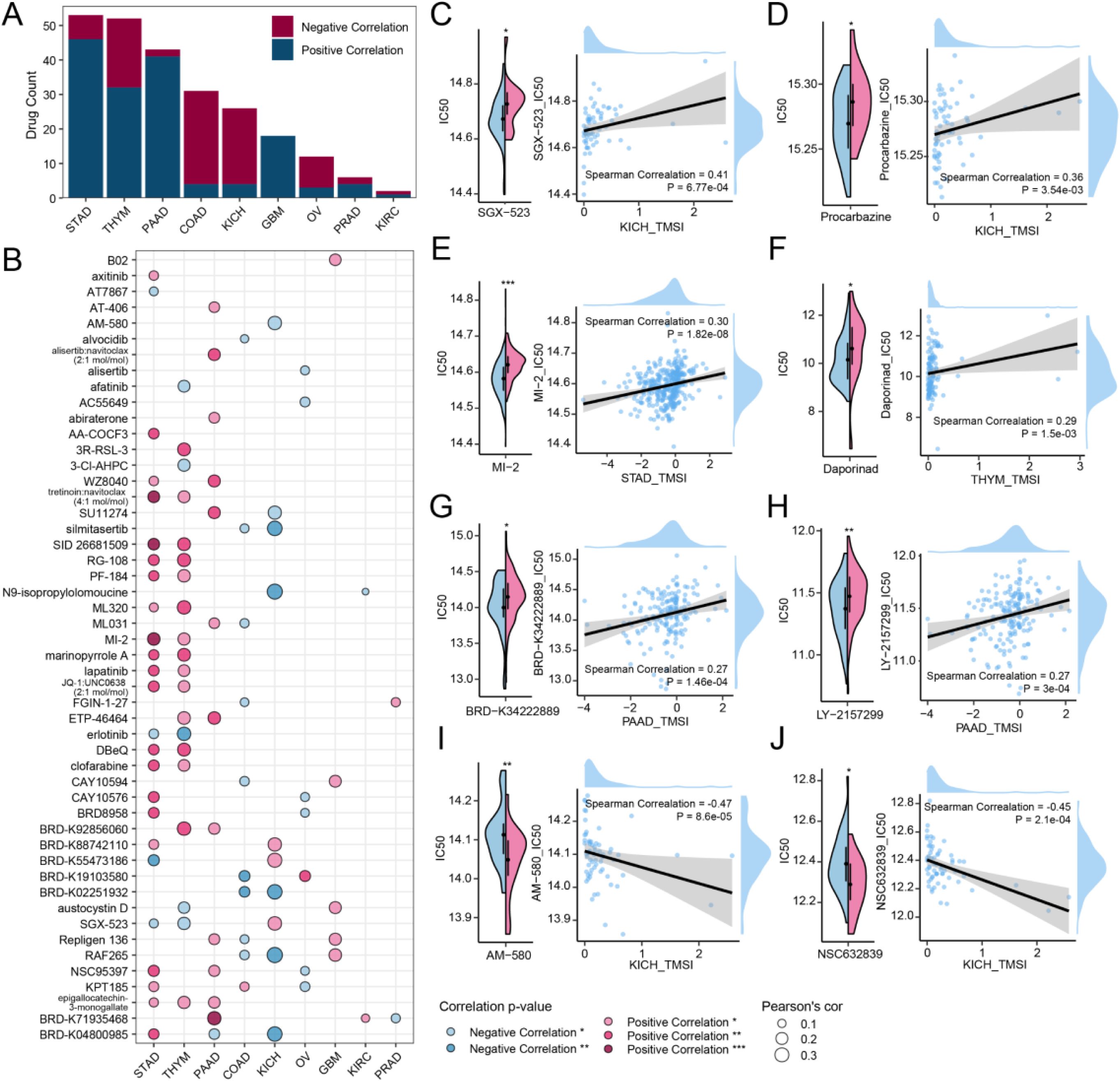
Evaluation of the predictive efficacy of TMSI models in chemotherapy. **A**. Statistics of drugs significantly different between the TMSI-high group and the TMSI-low group as well as correlated with the TMSI across 10 cancers. **B**. Presenting top 50 high-frequency drugs significantly correlated with TMSIs across each cancer. **C-J**. Violin plots and correlation plots of the comparison in IC50 values between the TMSI-high group and the TMSI-low group. The blue part represented the TMSI-low group, and the pink part represented the TMSI-high group in violin plots. **, p<0.01; *, p<0.05.

## Conclusion

In this study, we introduced the Tumor Microbiome Survival Index (TMSI), a model designed to evaluate the prognostic value of microbiome signatures for various cancer types. By analyzing microbiome profiles across ten different cancers, we identified specific tumor-associated microbes that are significantly correlated with patient survival.

Our approach goes beyond mere correlation, delving into the depths of biological mechanisms. We reveal the effectiveness of the TMSI model in predicting prognosis., proposing a tantalizing hypothesis where tumor-associated microbes could exert their influence by altering the gene methylation landscape, thereby modulating gene expression. As we stand on the cusp of a new frontier in cancer research, our findings suggest that the TMSI does not merely predict survival but also holds the key to unlocking the mysteries of chemotherapy sensitivity. Specifically, for patients afflicted with KICH, PAAD, STAD, or THYM, the TMSI emerges as a promising biomarker, offering a personalized therapeutic roadmap and a beacon of hope in the quest for precision medicine.

Although the TMSI model has demonstrated efficacy in predicting the prognosis of patients with various cancers, there are certain limitations that warrant further discussion. A major limitation is the lack of sufficient publicly available datasets to fully validate our findings; thus, we have only assessed the predictive capacity of TMSI for survival risk in patients with PAAD. Moreover, as our analysis relies on retrospective data, controlling for various potential confounding factors is challenging. Therefore, it is essential to design extensive prospective multicenter studies to ascertain the prognostic capabilities of tumor microbiome biomarkers. Additionally, while our research has begun to elucidate the interactions between tumor microbiome biomarkers and the tumor microenvironment and prognosis in different cancers, the underlying mechanisms still require further investigation.

In summary, our research offers new perspectives on the development of cancer and the intricate interactions between the immune system and microbial communities. Our joint analysis revealed potential interactions between TMSI microbiome signatures and tumor immune microenvironment, especially the humoral immune response. Moreover, TMSI has guiding significance in the use of chemotherapy treatments, potentially leading to more personalized therapeutic strategies for cancer patients.

## Acknowledgments

This work was partially supported by the National Key R&D Program of China (Grant No. 2023YFA1800900 and 2018YFC0910502), the National Natural Science Foundation of China (Grant Nos. 32071465, 31871334, 81827901). Numerical computations were performed on the Hefei Advanced Computing Center.

## Ethics approval and consent to participate

Not applicable.

